# Engineering a Matrix-Preserving Vascular dECM Platform with Tunable Stiffness for *In Vitro* Vascular Remodeling

**DOI:** 10.64898/2026.05.09.724001

**Authors:** Yuna Heo, Rhonda Drewes, Se-Hwan Lee, Yongho Bae, Su Chin Heo

## Abstract

Pathologic arterial stiffening is a hallmark of vascular disease that contributes to maladaptive vascular remodeling and neointimal hyperplasia through vascular smooth muscle cell (VSMC) phenotypic switching. Yet, because vascular disease progression is governed by both biomechanical and extracellular matrix (ECM) alterations, existing *in vitro* models often fail to recapitulate the full complexity of the diseased vascular microenvironment. Here, we developed a bioactive decellularized extracellular matrix (dECM) and methacrylated hyaluronic acid (MeHA) composite scaffold platform with tunable stiffness that preserves native vascular ECM components while enabling controlled investigation of stiffness-dependent cell behavior. Proteomic analyses confirmed retention of key vascular matrisome components, including collagens and glycoproteins, following decellularization. Electrospun vascular dECM scaffolds maintained an aligned fibrous architecture and spanned stiffness ranges representative of healthy and pathologically stiffened arterial microenvironments. Within this matrix-preserving platform, human VSMCs cultured on stiff dECM scaffolds exhibited increased spreading, altered morphology, enhanced nuclear localization of YAP and survivin, and broad transcriptional changes consistent with a shift toward a proliferative, matrix-remodeling VSMC phenotype. Together, this bioactive, matrix-preserving platform enables mechanobiologically relevant modeling of stiffness-driven vascular remodeling and indicates YAP and survivin as candidate regulators of maladaptive VSMC mechanotransduction.

## 1. INTRODUCTION

Cardiovascular disease (CVD) remains a leading cause of death in the United States and is estimated to affect more than 80 million people^1^. In 2022, nearly 20 million deaths worldwide were attributed to CVD, accounting for approximately 32% of all deaths^2^. The associated financial burden is substantial, with healthcare costs totaling approximately $168 billion from 2021 to 2022^3^. A defining feature of many vascular pathologies, including atherosclerosis, restenosis, and injury-associated neointimal hyperplasia, is progressive arterial wall stiffening. This stiffening arises from dynamic remodeling of extracellular matrix (ECM) composition and organization, creating an abnormal mechanical microenvironment for resident cells^4,5^. Within the tunica media, vascular smooth muscle cells (VSMCs) are embedded among collagen and elastic fibers, where they regulate vascular tone, maintain ECM homeostasis, and sense and respond to changes in matrix stiffness^6,7^. Under pathologic conditions, VSMCs undergo a phenotypic switch from a contractile to a synthetic state characterized by increased migration, proliferation, and ECM production^8,9^. These maladaptive responses contribute directly to neointimal hyperplasia/formation, luminal narrowing, and further ECM remodeling, positioning arterial stiffening as both a hallmark and an active driver of vascular disease progression (**Figure 1A**).

**Figure 1:**
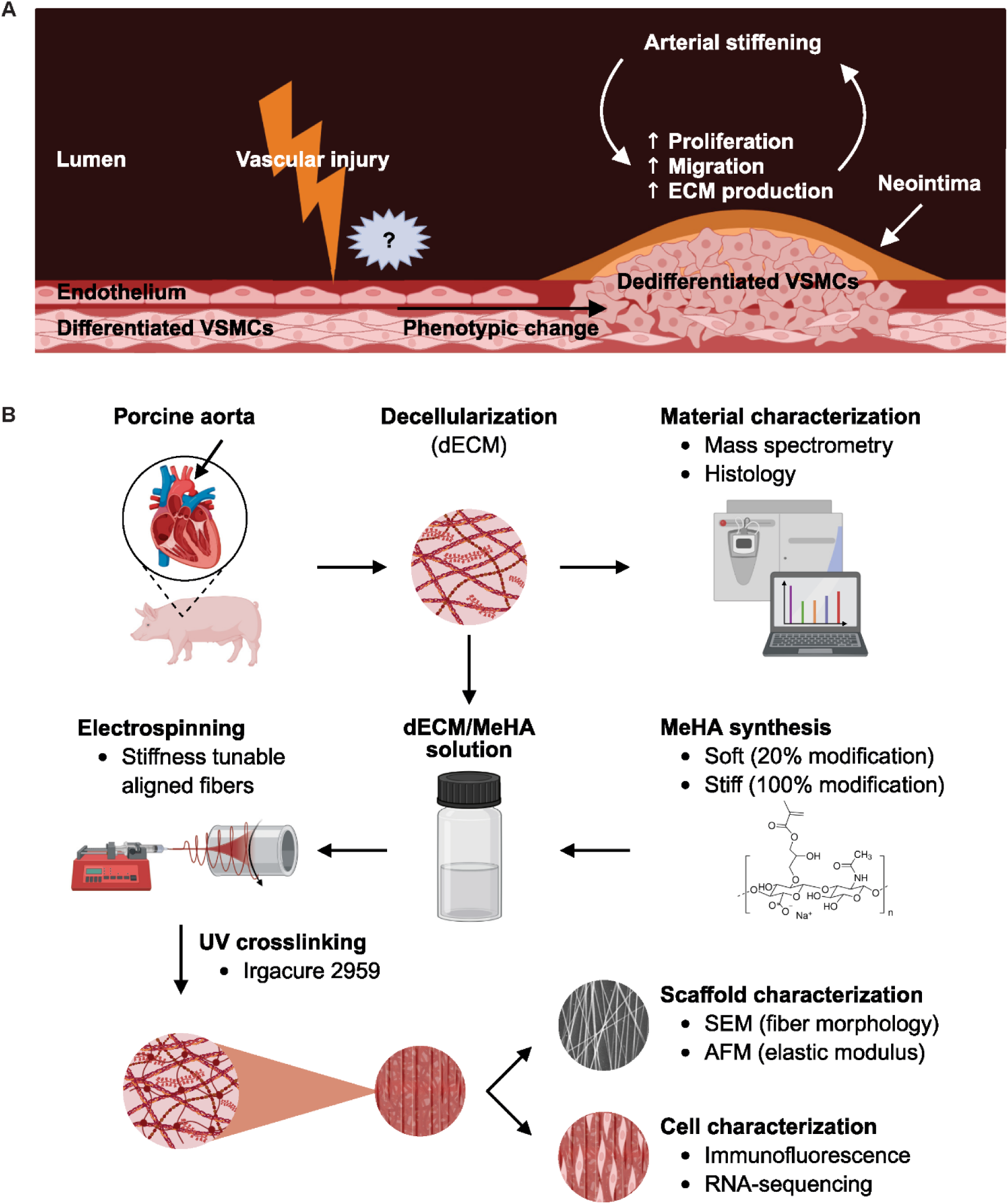
Arterial stiffening drives maladaptive VSMC remodeling and motivates the development of a stiffness-tunable vascular dECM scaffold platform. **(A)** Conceptual schematic illustrating vascular injury-associated arterial stiffening and its effects on vascular smooth muscle cell (VSMC) phenotypic switching, including increased proliferation, migration, extracellular matrix (ECM) production, and neointima hyperplasia. **(B)** Experimental workflow for porcine aorta decellularization, dECM characterization, MeHA synthesis, electrospinning-based fabrication of dECM–MeHA composite scaffolds, UV crosslinking, scaffold characterization, and downstream VSMC analyses. (Created with BioRender.com)

These pathologic VSMC changes are mediated in part by mechanotransduction pathways, which convert matrix-derived mechanical cues into biochemical signals that regulate transcriptional programs and cellular phenotype. In VSMCs, increased substrate stiffness has been shown to enhance cytoskeletal tension and alter nuclear morphology, thereby activating mechanosensitive transcriptional signaling pathways^10,11^. Among these, the Hippo pathway effector Yes-associated protein (YAP) is a central stiffness-responsive factor that shuttles between the cytoplasm and nucleus in response to cell shape, cytoskeletal organization, and matrix rigidity^12^. Nuclear YAP drives transcriptional programs associated with proliferation, survival, and ECM remodeling. Survivin (BIRC5), a pro-survival and cell-cycle regulator, has also emerged as a potential mechanosensitive mediator of vascular remodeling^13^. Prior studies have shown that survivin is upregulated following arterial injury and during neointimal formation, while inhibition reduces stiffness-mediated VSMC responses under pathological conditions^14–18^. Together, these findings suggest that YAP- and survivin-associated pathways contribute to stiffness-driven VSMC phenotypic switching.

Despite this recognition, there remains a need for *in vitro* models that recapitulate both the biochemical complexity of the arterial ECM and the pathologic stiffness of diseased vessels. Conventional mechanobiology platforms, including polyacrylamide and other synthetic hydrogels, enable precise control over substrate stiffness but lack the biochemical complexity, fibrous architecture, and tissue-specific composition of native vascular ECM^19,20^. Conversely, decellularized ECM (dECM) platforms preserve tissue-specific biochemical cues but are often implemented as coatings, hydrogels, or bulk tissues with limited control over stiffness^21,22^. Consequently, it remains difficult to model how progressive arterial stiffening regulates VSMC behavior within a preserved vascular ECM context, particularly across the transition from healthy to pathologically stiffened microenvironments.

To address this unmet need, we engineered a stiffness-tunable *in vitro* model of the vascular microenvironment by integrating porcine aorta-derived dECM with methacrylated hyaluronic acid (MeHA) in an electrospun scaffold platform (**Figure 1B**). This design preserved key vascular ECM components and fibrous architecture while enabling controlled modulation of substrate stiffness across physiologic and pathologic ranges. We hypothesized that pathologic stiffness within this vascular dECM platform promotes maladaptive VSMC behavior, including increased cell spreading, enhanced nuclear localization of YAP and survivin, and transcriptional programs associated with a synthetic and disease-associated phenotype relative to soft scaffolds. This study establishes a biomimetic *in vitro* platform for investigating stiffness-mediated vascular degeneration and provides a translational framework for dissecting mechanobiological pathways involved in neointimal formation, vascular remodeling, and vascular arterial disease progression.

## 2. RESULTS

### 2.1. Decellularization preserves core vascular matrisome components required for microenvironment fidelity

To establish an *in vitro* model capable of recapitulating vascular microenvironments, we first evaluated the preservation of native vascular ECM composition following decellularization. Proteomic comparison of native and decellularized (dECM) porcine aorta confirmed effective cell removal while largely preserving the structural matrix composition relevant for vascular microenvironment modeling. Principal component analysis (PCA) demonstrated clear separation between native and dECM samples along PC1 (**Supplementary Figure 1A**), and heatmap analysis of selected proteins highlighted these compositional differences (**Supplementary Figure 1B**). Importantly, across all samples, 4,737 proteins were detected, of which the majority (4,389; ∼93%) were shared between native and dECM tissues, indicating that the decellularization process conserves most of the vascular proteome (**Figure 2A**). A subset of proteins was unique to native tissue (343; ∼7%), whereas only 5 proteins were exclusively detected in dECM, suggesting that decellularization primarily removed a limited subset of native proteins while introducing minimal processing-related artifacts.

**Figure 2:**
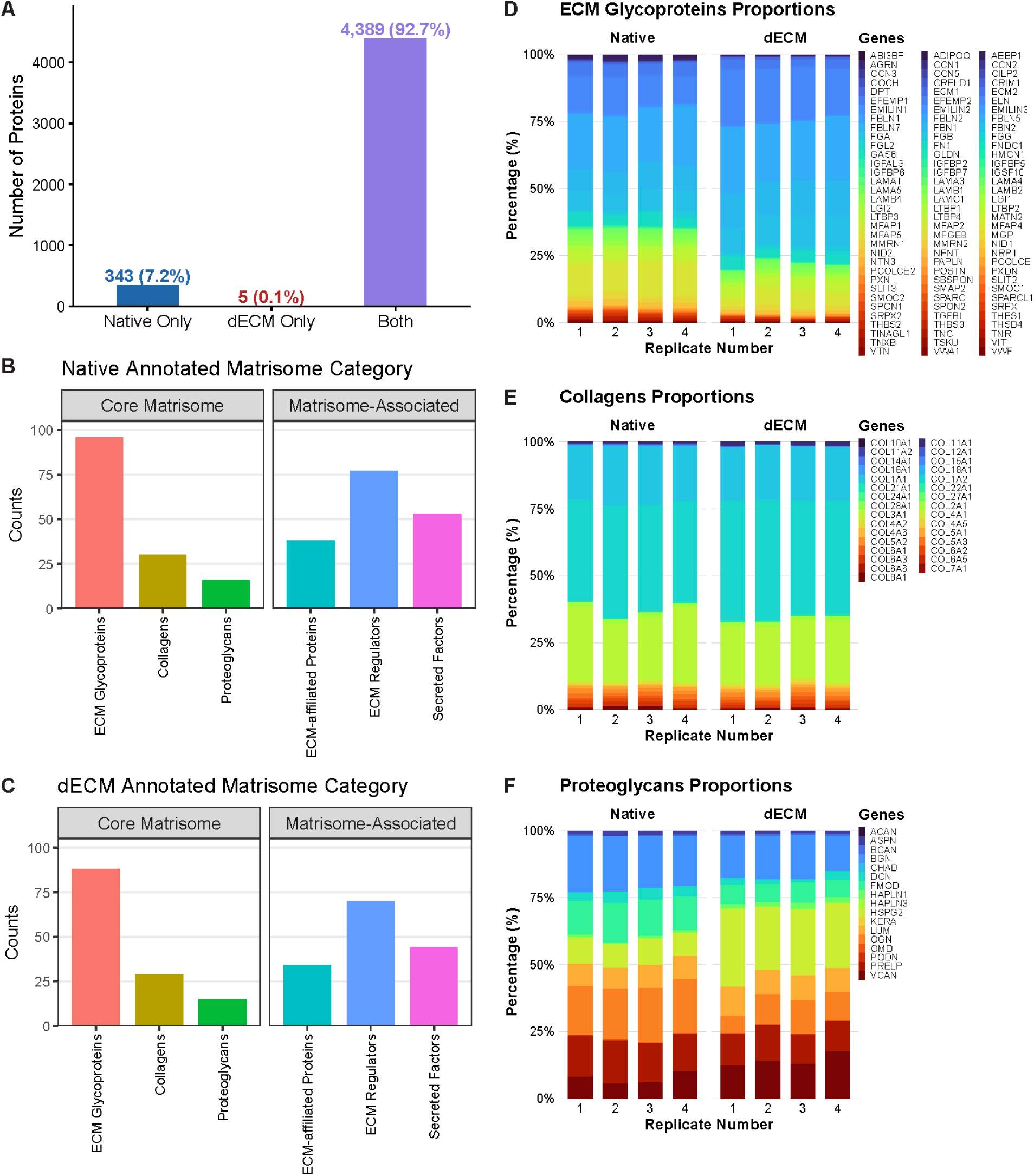
Proteomic analysis demonstrates broad retention of vascular ECM proteins following porcine aorta decellularization. **(A)** Overlap of proteins identified in native and decellularized (dECM) porcine aorta samples, showing proteins uniquely detected in native tissue, uniquely detected in dECM, or shared between both groups. Matrisome annotation of proteins detected in **(B)** native and **(C)** dECM samples, categorized into core matrisome and matrisome-associated classes. Relative distribution of core matrisome subclasses across native and dECM samples, including **(D)** ECM glycoproteins, **(E)** collagens, and **(F)** proteoglycans (n = 4 biological replicates per group).

Because vascular degeneration is fundamentally associated with ECM remodeling, retention of core matrisome components was specifically examined to determine whether dECM preserved the biochemical fidelity required to model the vascular microenvironment. Matrisome annotation revealed that core structural ECM classes, including collagens, ECM glycoproteins, and proteoglycans, were retained at similar counts and relative proportions in native and dECM samples (**Figure 2B-F**). This conservation indicates preservation of fibrillar and pericellular matrix components after decellularization, which is critical for maintaining the biochemical and structural context through which VSMCs sense and respond to pathologic mechanical cues. In contrast, matrisome-associated proteins (ECM-affiliated proteins, ECM regulators, and secreted factors) exhibited greater variability between native and dECM (**Supplementary Figure 2A-C**), consistent with the expected depletion of soluble factors and regulatory proteins during detergent-based decellularization. These findings indicate that the decellularization process preferentially preserved structural matrix identity while reducing the content of regulatory proteins.

Gene ontology enrichment analysis further supported retention of vascular matrix-associated functional annotations after decellularization, with dECM-enriched terms including extracellular matrix organization, collagen fibril organization, and cell adhesion. (**Supplementary Figure 1C**). These enriched terms highlight preservation of structural and adhesive functions necessary for physiologically relevant VSMC-ECM interactions. Histological analyses corroborated these proteomic findings: hematoxylin and eosin (H&E) staining confirmed effective removal of cellular nuclei and cytoplasmic components, while picrosirius red (PSR) staining demonstrated preservation of collagen architecture in dECM relative to native tissue (**Supplementary Figure 3**).

Together, these data demonstrate that porcine aorta decellularization effectively removed cellular components while preserving the core vascular matrisome and structural framework, thereby establishing a matrix-preserving foundation for engineering an *in vitro* stiffness-tunable vascular microenvironment model.

### 2.2. Vascular dECM-based nanofibrous scaffolds provide a matrix-preserving platform with tunable stiffness

To model physiologic and pathologic arterial mechanics within a matrix-preserving context, we blended vascular dECM powder with methacrylated hyaluronic acid (MeHA) of low (“soft”, ∼25% methacrylation) or high (“stiff”, ∼100% methacrylation) modification (**Supplementary Figure 4**) and electrospun into aligned nanofibrous scaffolds. This composite design enabled mechanical tunability while retaining vascular-derived matrix components within a fibrous architecture. Scanning electron microscopy (SEM) revealed dense and aligned fiber networks in both soft and stiff scaffolds, with comparable fiber diameters between groups (**Figure 3A-B**). These data indicate that variations in MeHA methacrylation and crosslinking primarily modulated mechanical properties without significantly disrupting scaffold structure.

**Figure 3:**
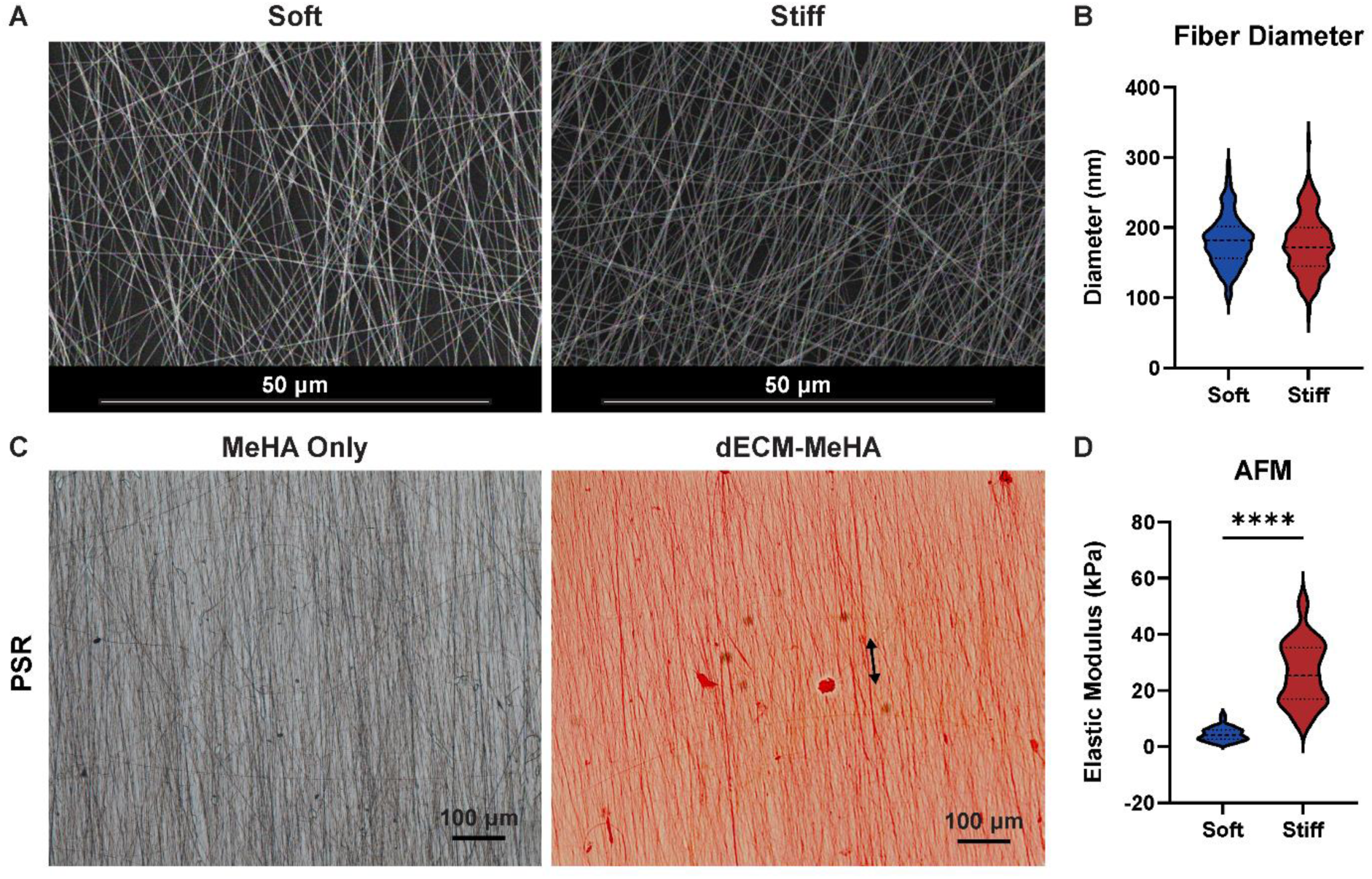
MeHA incorporation enables stiffness modulation of vascular dECM-based nanofibrous scaffolds while preserving fibrous scaffold morphology and matrix incorporation. **(A)** Representative scanning electron microscopy (SEM) images of soft and stiff dECM-MeHA electrospun scaffolds demonstrating preserved nanofibrous architecture across stiffness conditions (scale bars = 50 µm). **(B)** Quantification of fiber diameter in soft and stiff scaffolds (n > 100 fibers per group; median with interquartile range). **(C)** Representative picrosirius red (PSR) staining of MeHA-only and dECM-MeHA scaffolds demonstrating collagen-containing matrix incorporation within dECM–MeHA scaffolds (scale bars = 100 µm). **(D)** Atomic force microscopy (AFM)-measured elastic modulus of soft and stiff scaffolds (soft: n > 100; stiff: n = 39 measurements; median with interquartile range; unpaired t-test; ****p < 0.0001).

To assess ECM incorporation, PSR staining confirmed robust collagen signal in dECM-MeHA scaffolds, whereas MeHA-only control showed minimal staining (**Figure 3C**). These results demonstrate successful integration of vascular ECM components into the composite scaffold system, establishing a collagen-rich, matrix-associated biochemically complex environment, whereas MeHA-only scaffolds primarily provide a synthetic structural backbone.

Mechanical characterization by atomic force microscopy (AFM) indentation revealed a distinct separation in elastic modulus between soft and stiff dECM-MeHA scaffolds (**Figure 3D**). Soft scaffolds exhibited a low-kPa stiffness range (∼5 kPa), consistent with more compliant vascular tissue. In contrast, stiff scaffolds reached a higher range (∼26 kPa), which approximates the pathologic arterial stiffness observed in atherosclerosis and related vascular pathologies^23^. Importantly, this stiffness modulation was achieved without significant differences in fiber morphology and ECM incorporation, allowing subsequent VSMC responses to be attributed primarily to mechanical cues rather than structural or compositional differences.

Together, these results establish vascular dECM-based nanofibrous scaffolds as a matrix-preserving, stiffness-tunable platform for controlled investigation of mechanical regulation of cell behavior within a conserved ECM environment.

### 2.3. Stiffness-dependent changes in VSMC morphology, YAP activation, and survivin localization

To determine how scaffold stiffness influences VSMC behavior within a matrix-preserving platform, we evaluated stiffness-dependent changes in cell morphology and mechanosensitive signaling at the single-cell level. After 24 hours of culture, VSMCs on soft scaffolds exhibited smaller, more rounded morphologies, whereas cells on stiff scaffolds were more spread and elongated (**Figure 4A**). Quantitative image analysis confirmed that VSMCs on stiff scaffolds exhibited significantly increased cell area, reduced roundness, and increased aspect ratio relative to cells on soft scaffolds (**Figure 4B-D**). These findings indicate that increased scaffold stiffness promotes pronounced cytoskeletal remodeling, consistent with a mechanically activated cellular state^24,25^.

**Figure 4:**
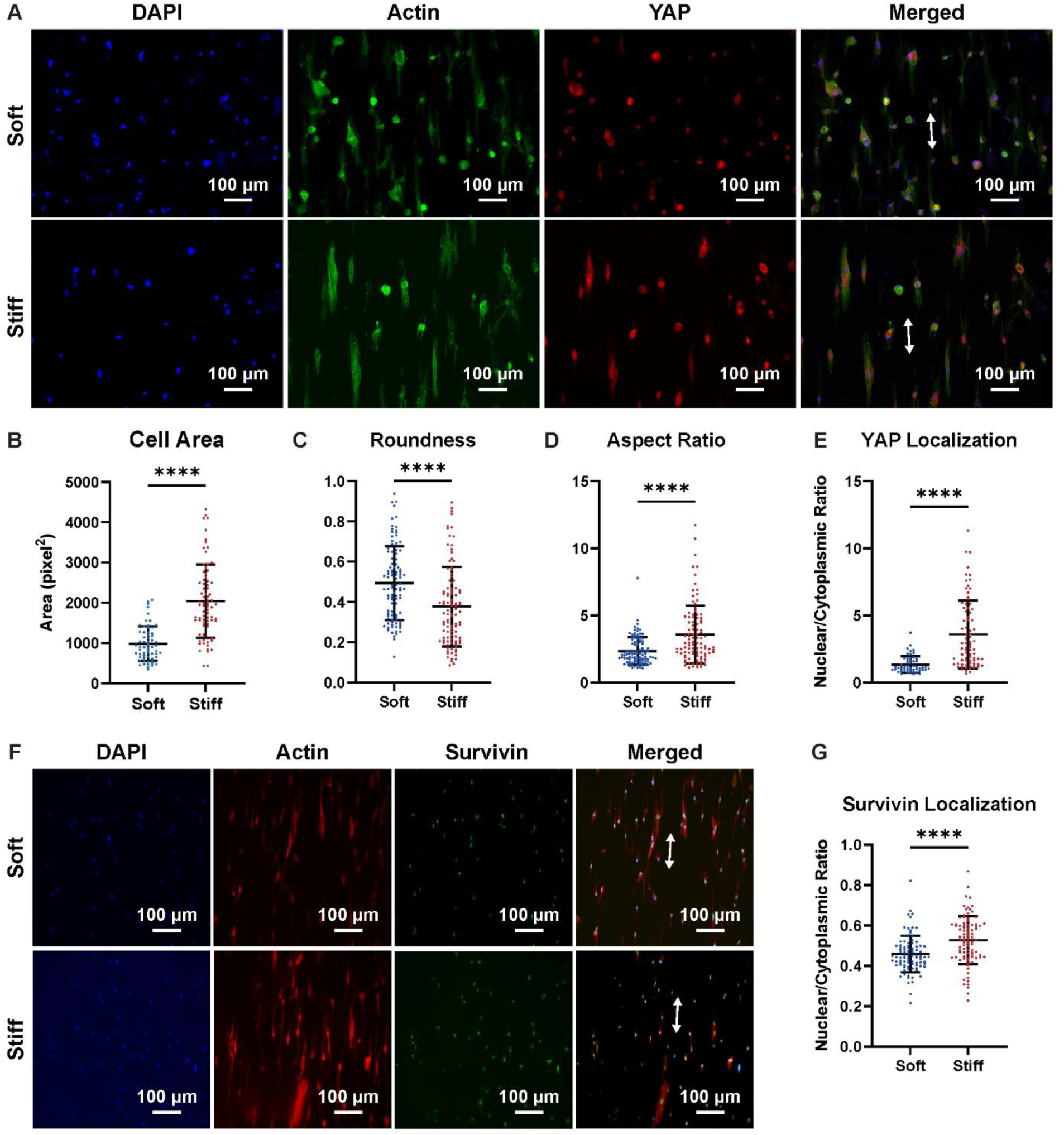
Increased scaffold stiffness promotes VSMC morphological remodeling, YAP nuclear localization, and survivin nuclear localization within vascular dECM-based scaffolds. **(A)** Representative immunofluorescence images of VSMCS cultured on soft and stiff scaffolds and stained for DAPI, actin, and YAP (scale bars = 100 µm). Single-cell quantification of VSMC morphology, including **(B)** cell area, **(C)** roundness, and (D) aspect ratio (n > 90 cells; mean ± SD; unpaired t-test; **** p < 0.0001). (E) Quantification of YAP nuclear-to-cytoplasmic fluorescence intensity ratio in VSMCs cultured on soft and stiff scaffolds. **(F)** Representative immunofluorescence images stained for DAPI, actin, and survivin (scale bars = 100 µm). (G) Quantification of nuclear-to-cytoplasmic fluorescence intensity ratio in VSMCs cultured on soft and stiff scaffolds (n > 90 cells; mean ± SD; unpaired t-test; **** p < 0.0001).

Given that YAP is a key mechanosensitive transcriptional regulator, we next examined its subcellular localization response to scaffold stiffness^26^. Immunofluorescence imaging demonstrated increased nuclear YAP accumulation in VSMCs cultured on stiff scaffolds compared with soft scaffolds. Quantification of the nuclear-to-cytoplasmic YAP ratio confirmed significantly elevated nuclear localization under stiff conditions (**Figure 4E**). These observations suggest that increased scaffold stiffness within a preserved vascular ECM context promotes activation of YAP-dependent mechanotransduction. Because scaffold architecture and matrix composition were largely conserved across conditions, these findings support mechanical stiffness as a primary contributor to VSMC mechanobiological activation/reprogramming.

Given the established role of survivin (BIRC5) in cell-cycle regulation, proliferation, and vascular ECM remodeling, as well as its regulation by mechanical cues^14,15^, we next examined its expression and subcellular localization as a potential downstream readout of stiffness-associated phenotypic adaptation. On soft scaffolds, survivin was distributed across cytoplasmic and nuclear compartments, whereas VSMCs cultured on stiff scaffolds exhibited more pronounced nuclear enrichment of survivin (**Figure 4F**), paralleling the stiffness-dependent nuclear localization observed for YAP. Quantification of the nuclear-to-cytoplasmic survivin ratio confirmed a significant increase under stiff conditions (**Figure 4G**), identifying that matrix stiffening promotes nuclear survivin localization and signaling, which may support enhanced cell-cycle progression and adaptive ECM remodeling.

Together, these findings demonstrate that increased stiffness within vascular dECM-based scaffolds drives coordinated VSMC morphological remodeling and promotes nuclear localization/activation of YAP and survivin. This mechanobiological response supports the utility of this platform for modeling stiffness-mediated vascular cell reprogramming.

### 2.4. Stiffness-induced transcriptomic remodeling reveals phenotypic reprogramming in VSMCs

To determine whether stiffness-dependent morphological and mechanotransductive changes were associated with broader phenotypic reprogramming, we performed RNA-sequencing on VSMCs from two donors cultured on soft or stiff scaffolds for 24 hours. PCA demonstrated partial separation between conditions (**Supplementary Figure 5A**), indicating stiffness-dependent transcriptional divergence.

Across donors, stiff scaffolds induced broad changes in gene programs associated with cell proliferation, ECM remodeling, and VSMC phenotypic regulation (**Figure 5A-D**). Expression of multiple cell-cycle-associated genes, including *CCND1*, *CCNA1*, *CCNA2*, *CCNB1*, *CCNB2*, *E2F1*, and *BIRC5* (survivin), was increased under stiff conditions (**Figure 5A**), consistent with a proliferative phenotype^14,27^. In parallel, expression of matrix-related genes was elevated (**Figure 5B**), suggesting enhanced ECM remodeling in response to increased substrate stiffness. Notably, contractile markers, including *MYH11*, *LMOD1*, *CALD1*, *SMTN*, *ACTA2*, *TAGLN*, and *CNN1*, were maintained or modestly increased under stiff conditions (**Figure 5C**), indicating that stiffness does not uniformly suppress contractile gene expression. Concurrently, several synthetic markers, including *SPP1*, *FBP1*, *LGALS3*, *VIM*, *KLF4*, *MYH10*, and *TPM4*, were also upregulated (**Figure 5D**), reflecting activation of synthetic features. Together, these data indicate that stiff dECM scaffolds induce a complex VSMC response characterized by proliferative activation, ECM remodeling, and changes in both contractile- and synthetic-associated genes, rather than a binary contractile-to-synthetic switch.

**Figure 5:**
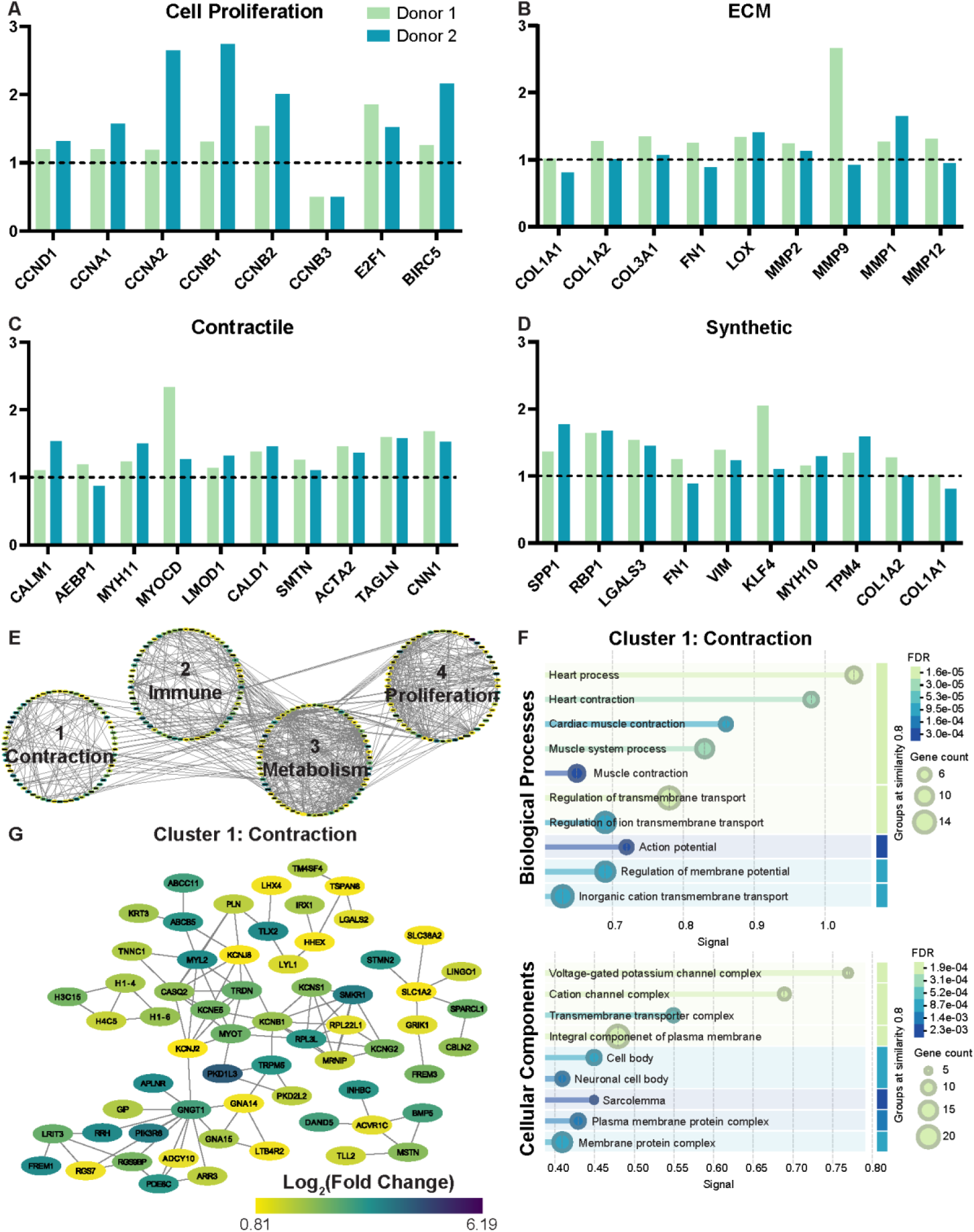
RNA-sequencing reveals stiffness-responsive transcriptional programs associated with proliferation, ECM remodeling, VSMC phenotype regulation, and contraction-associated interaction networks. Relative gene expression changes in VSMCs cultured on stiff versus soft dECM-MeHA scaffolds for 24 hours from two independent biological donors (Donor 1 and 2). Gene sets include markers associated with **(A)** cell proliferation, **(B)** ECM remodeling, **(C)** contractile VSMC phenotype, and **(D)** synthetic VSMC phenotype. Dashed lines at 1 indicate no change between stiff and soft conditions. **(E)** STRING-based protein–protein interaction network and clustering analysis of significantly upregulated differentially expressed genes (DEGs), identifying major functional clusters associated with contraction, immune-related signaling, metabolism/stress responses, and proliferation. **(F)** STRING interaction network of the contraction-associated cluster, with node color indicating log2(fold change). **(G)** Gene ontology enrichment analysis of the contraction-associated cluster, highlighting enriched biological process and cellular component categories. RNA-sequencing was performed using VSMCs from n = 2 biological donors cultured on soft and stiff scaffolds for 24 hours. Differential expression analysis was performed using DESeq2, with significance defined as p < 0.5 and fold change > 2.0.

To further define stiffness-responsive biological programs, we performed STRING-based clustering and protein-protein interaction (PPI) analysis and identified four major transcriptional clusters (**Figure 5E, Supplementary Figure 5B-D**). GO enrichment analysis of cluster 1 identified contraction-associated biological processes, including heart contraction, cardiac muscle contraction, and muscle contraction (**Figure 5F**), suggesting engagement of cytoskeletal and force-regulatory programs in VSMCs on stiff scaffolds. PPI analysis identified highly interconnected genes within this cluster, including *MYL2*, which regulates myosin motor activity, and *TRDN*, a calcium regulator of cardiac muscle contraction (**Figure 5G**). These findings are consistent with the established role of VSMCs in maintaining vascular tone through modulated contraction and control of vessel diameter, processes essential for proper blood flow and lumen homeostasis^28,29^. Cluster 2 demonstrated enrichment of immune-related processes, including adaptive immune response and immune system process (**Supplementary Figure 5B, 6A**), consistent with emerging roles of VSMCs in inflammatory signaling during vascular disease^30^. PPI analysis identified highly interconnected genes within this cluster, including *GZMK*, associated with pro-inflammatory activity^31,32^, and *CRTAM*, a modulator of immune responses^33,34^. Cluster 3 was enriched for metabolism- and stress-related GO terms, including responses to lipids, bacteria, and SMAD signaling (**Supplementary Figure 5C, 6B**), indicating broader adaptive transcriptional responses to stiffness. Key interconnected genes included *JUN*, *FOS*, *CXCL8*, *PTGS2*, and *CEBPB*, suggesting the activation of metabolic and stress-responsive transcriptional programs in VSMCs beyond proliferative pathways. Cluster 4 was enriched for cell-cycle-related pathways, including nuclear chromosome segregation and nuclear division (**Supplementary Figure 5D, 6C**). Highly interconnected regulators included *PTTG1*, *SPC25*, and *TTK*, which are involved in sister chromatid separation, kinetochore function, and mitotic checkpoint regulation^35–37^, respectively, indicating robust activation of proliferative machinery. This program is consistent with VSMCs phenotypic transitions observed in atherosclerosis, where increased proliferation contributes to plaque development and stiffening. Together, these transcriptomic data suggest that increased scaffold stiffness promotes coordinated VSMC transcriptional programs associated with proliferation, ECM remodeling, and phenotypic adaptation, supporting this platform as a useful model for studying stiffness-driven vascular remodeling.

## 3. DISCUSSION

Arterial stiffening is a defining feature of vascular disease and contributes to maladaptive VSMC remodeling; however, many existing *in vitro* systems used to study stiffness-mediated VSMC behavior often fail to combine controlled stiffness modulation with preservation of the biochemical and structural complexity of native vascular ECM. In this study, we developed a matrix-preserving, stiffness-tunable vascular dECM platform that addresses this gap by integrating vascular ECM complexity with controlled modulation of substrate stiffness. Using this system, we found that increased stiffness promoted VSMC spreading, nuclear localization of YAP and survivin, and transcriptional programs associated with cell proliferation, ECM remodeling, and phenotypic adaptation. Together, these findings suggest that this platform provides a useful *in vitro* model for studying stiffness-mediated VSMC function and remodeling within a biologically relevant vascular microenvironment.

A major goal of this work was to establish an *in vitro* model that better captures both the mechanical and biochemical features of the vascular ECM. Previous studies using synthetic hydrogels have provided important mechanistic insight into stiffness sensing, but lack tissue-specific ECM composition and architecture^38^. Conversely, decellularized ECM platforms preserve biochemical complexity but often provide limited control over mechanical properties. By combining porcine aorta-derived dECM with MeHA and processing the composite into electrospun nanofibers, this study establishes a matrix-rich fibrous platform in which arterial-like stiffness can be examined within a vascular ECM context. Preservation of core matrisome components after decellularization supports the biological relevance of this platform, while stiffness tunability enables controlled investigation of matrix-derived mechanical cues. This is particularly important because VSMCs *in vivo* respond not only to stiffness, but also to ECM ligands, fiber architecture, and matrix-derived signals^10,25^.

A central finding of this study is that increased stiffness within a matrix-preserving platform promoted nuclear localization of both YAP and survivin in VSMCs. YAP is a well-established mechanosensitive transcriptional regulator that responds to matrix stiffness, cytoskeletal tension, and cell shape^39,40^. In our system, VSMCs cultured on stiff substrates were more spread and elongated, consistent with increased cytoskeletal tension and mechanotransduction. The corresponding increase in nuclear YAP suggests that the dECM-based platform recapitulates stiffness-sensitive signaling observed in other mechanobiology systems, while providing a more biologically relevant vascular ECM context. Because YAP regulates genes involved in proliferation and survival^41–43^, its activation may represent an important mechanistic link between arterial stiffening and the transcriptional reprogramming of VSMCs during vascular disease.

Survivin may serve as a complementary mechanosensitive regulator linking matrix stiffness to VSMC proliferation. While survivin is classically recognized for its role in mitosis, cytokinesis, and cell survival^47,48^, it has also been implicated in vascular injury, atherosclerosis, hypertension, and stiffness-mediated VSMC behavior^14–17,44^. In this study, increased stiffness promoted nuclear survivin localization, a pattern previously associated with proliferative activity in cancer cells^13,45^, accompanied by upregulation of cell-cycle-related genes, including cyclins. These findings support a model in which matrix stiffening promotes a proliferative VSMC state through survivin-associated signaling^14^. Importantly, the concurrent activation of YAP and survivin raises the possibility that these pathways may act in a coordinated manner during stiffness-induced VSMC proliferation and vascular remodeling. Although this study does not establish a direct causal relationship between YAP and survivin, both emerge as candidate mediators of stiffness-dependent phenotypic adaptation.

The transcriptomic response to increased stiffness further suggests that VSMC adaptation is not defined by a simple binary switch from a contractile to synthetic phenotype. Instead, VSMCs displayed a broader remodeling state characterized by increased expression of proliferation-associated genes, ECM remodeling genes, and several synthetic markers, while also maintaining or modestly increasing contractile-associated genes. These findings suggest that stiffness-induced VSMC responses represent a multidimensional adaptive state rather than a singular phenotypic change. In the context of arterial stiffening, VSMCs may initially retain elements of their contractile identity while acquiring proliferative, inflammatory, or matrix-remodeling features that contribute to disease progression. The aligned fibrous architecture and vascular dECM composition of this platform may also help preserve aspects of contractile-associated gene expression, even under stiff conditions. The enrichment of broader terms, including immune-, metabolic-, and contraction-associated clusters in stiffness-induced remodeling, is further highlighted in this disease modeling platform. In atherosclerosis, VSMCs can adopt a macrophage-like phenotype in a chronic proinflammatory state, which contributes to disease progression and arterial stiffening^46,47^. Also within these disease states, VSMC metabolism may exhibit flexibility and is known to reprogram, interwoven with inflammatory responses^48^. This increased mechanical load in matrix-enriched areas has also been shown to dysregulate hemodynamics in vessels and further complicate aortic wall integrity and reduce heart function^49^. These findings further suggest that stiffness-induced remodeling extends beyond proliferation alone and may prime broader maladaptive states relevant to vascular disease.

While these results support the utility of this platform for modeling stiffness-dependent vascular remodeling, several limitations should be considered. First, although dECM preserves tissue-specific biochemical complexity, decellularization and scaffold fabrication likely alter the native organization, abundance, and accessibility of certain ECM components, including structural collagens, elastin-associated proteins, proteoglycans, and matricellular proteins. Additional biochemical and structural characterization would further clarify how closely this platform recapitulates native arterial ECM. Second, although nuclear localization of YAP and survivin suggests activation of mechanosensitive and proliferative pathways, functional studies are needed to determine whether these pathways directly regulate VSMC proliferation, migration, or matrix synthesis. Third, this platform does not yet incorporate endothelial interactions, immune cell components, or hemodynamic forces, all of which contribute to vascular disease progression *in vivo*. Finally, RNA-sequencing was performed using two human donors and demonstrated some donor-dependent variability. Although statistical significance was not achieved due to the limited donor samples, we have identified several observed trends from our analysis. Additional donor samples will be important for defining conserved stiffness-responsive pathways and distinguishing them from donor-specific responses.

Overall, this work establishes a matrix-preserving, stiffness-tunable vascular dECM platform for studying VSMC mechanobiology in a biologically relevant vascular ECM environment. These findings demonstrate increased stiffness promoted morphologic remodeling, YAP and survivin nuclear localization, and broad transcriptional changes associated with proliferation, ECM remodeling, and phenotypic plasticity. Together, these findings support a model in which arterial stiffening contributes to vascular remodeling by activating mechanosensitive pathways that drive VSMC adaptation and reprogramming. This platform may therefore provide a valuable foundation for future mechanistic studies and therapeutic screening aimed at targeting stiffness-driven pathways in vascular disease.

## 4. MATERIALS AND METHODS

### 4.1. Decellularized porcine aorta ECM preparation

Adult porcine aortas (n=4 biological replicates, Animal Technologies, Inc.) were decellularized based on previously reported protocols with adjustments^50^. Briefly, aortic tissues were finely minced into approximately 2 mm³ pieces and washed in deionized water for 6-9 hours under gentle agitation at room temperature. Subsequently, samples were then treated with 0.3% (w/v) sodium dodecyl sulfate (SDS; Thermo Fisher Scientific, 15525017) in phosphate-buffered saline (PBS; GrowCells, MRGF-6235) for 24 hours, followed by 3% (v/v) Triton-X100 (Sigma-Aldrich, X100) in PBS for 24 hours. To remove residual nucleic acids, tissues were incubated in 15 U/ml deoxyribonuclease (DNase; Sigma-Aldrich, D5025) supplemented with 50 mM MgCl_2_ (Sigma-Aldrich, 208337) in PBS for 24 hours at 37°C. Following each treatment, samples were washed thoroughly with PBS. Decellularized tissues were sterilized in 0.1% (v/v) peracetic acid (Sigma-Aldrich, 269336) and 4% (v/v) ethanol in PBS for 4 hours, then extensively rinsed, lyophilized, and cryo-milled into a fine powder for downstream scaffold fabrication.

### 4.2. Proteomic analysis

To evaluate protein preservation following decellularization, lyophilized native and decellularized tissues (n=4 biological replicates per group) were cryo-milled and processed for liquid chromatography-tandem mass spectrometry (LC-MS/MS) analysis as previously described^57^. Proteins identified in at least 2 of 4 biological replicates within each group were included for downstream analysis. Differential protein abundance between native and decellularized tissues was assessed using the limma package in R, and multiple hypothesis testing was corrected using the Benjamini-Hochberg false discovery rate (FDR) correction^51^. Proteins with p-value < 0.05 were considered significantly differentially abundant. Matrisome annotation and category classification were performed using MatrisomeAnalyzeR^52^. Gene ontology (GO) enrichment analyses were conducted and visualized using the ggplot2 package in R.

### 4.3 MeHA synthesis

Methacrylated hyaluronic acid (MeHA) was synthesized according to previously established protocols^50,53^. Sodium hyaluronate (Lifecore) was dissolved 1% (w/v) in deionized water overnight. Methacrylic anhydride (Sigma-Aldrich, 276685) was added dropwise incrementally, while maintaining the solution pH at 8.0-8.5 using 5N NaOH (Fisher Scientific, SS2551). The reaction was carried out on ice with continuous stirring to control reaction kinetics and minimize degradation. The degree of methacrylation was modulated by varying the amount of methacrylic anhydride added, targeting approximately 20% modification (for “soft”) and approximately 100% modification (for “stiff”). Following synthesis, MeHA solutions were dialyzed against deionized water to remove unreacted reagents, then frozen and lyophilized. The degree of methacrylation was quantified by ^1^H-NMR (10 mg/mL in D_2_O; Bruker NEO400) (**Supplementary Figure 4**). Degree of methacrylation was calculated from the ratios of the areas under the characteristic methacrylate proton peaks at 5.6 and 6.1 ppm to the peak at 1.9 ppm (N-acetyl glucosamine of HA)^54–57^.

### 4.4. dECM-based stiffness-tunable nanofibrous scaffold fabrication

Decellularized ECM (dECM) powder was solubilized in an acetic acid (Sigma-Aldrich, 1.00066), ethyl acetate (Sigma-Aldrich, 270989), and deionized water solvent system (AED; 3:2:1, v/v/v). Briefly, 1.4 g of dECM powder was dissolved in 10 mL of AED solution and mixed at 45°C until fully solubilized. The resulting dECM solution was then combined with 4% (w/v) MeHA (“soft” or “stiff”), 2% (w/v) poly(ethylene oxide) (PEO; Thermo Fisher Scientific, 183221000), and 0.5% (w/v) Irgacure (Sigma-Aldrich, 410896) to generate electrospinning precursor solutions. The prepared solutions were electrospun using an 18-gauge needle (Hamilton) at a flow rate of 1 mL/h. Fibers were collected onto a grounded rotating mandrel at a 15 cm tip-to-collector distance with an applied voltage of 17 kV. Electrospinning was performed under controlled temperature and humidity conditions. Following fabrication, the electrospun scaffolds were photo-crosslinked under UV light for 30 minutes to stabilize scaffold structure and establish stiffness-dependent mechanical properties.

### 4.5. Scaffold characterization

Scaffold fiber morphology was examined by scanning electron microscopy (SEM). Electrospun scaffolds were sputter-coated with a gold-palladium alloy and imaged on a FEI Quanta 250 FEG scanning electron microscope (Thermo Fisher Scientific) under high-vacuum conditions at an accelerating voltage of 10 kV. Fiber diameter was quantified from SEM images using ImageJ (NIH).

Atomic force microscopy (AFM) was used to quantify scaffold stiffness. Scaffold surfaces were indented using a silicon nitride cantilever with a three-sided pyramidal tip (Cat. No. BL-AC40TS-C2; Asylum Research; spring constant, 0.09 N/m; tip radius, 8 nm). Measurements were acquired in contact mode using an NX12 AFM system (Park Systems) mounted on a Nikon ECLIPSE Ti2 inverted microscope. Before measurement, scaffolds were hydrated in PBS for at least 20 minutes. Approximately 10-20 measurements were recorded per sample. Force-distance curves were analyzed using XEI software (Park Systems) and converted to stiffness measurements.

To assess collagen preservation of fabricated scaffold, MeHA-only and dECM-MeHA scaffolds were stained with picrosirius red (PSR; picric acid, Thermo Fisher Scientific, 18612375; Direct Red 80, Sigma-Aldrich, 365548) and imaged using Axioscanner (Zeiss). For histologic comparison of tissue structure before and after decellularization, native and decellularized porcine aorta samples were stained with hematoxylin and eosin (H&E; hematoxylin solution, Sigma-Aldrich, GHS232; eosin Y solution, Sigma-Aldrich, HT110132) and PSR staining.

### 4.6. Cell culture

Human vascular smooth muscle cells (VSMCs; Cell Applications, Inc., 354-05a) were cultured in Dulbecco’s modified Eagle’s medium (DMEM; GIBCO, 11885084) supplemented with 10% fetal bovine serum (FBS; GIBCO, 2510268RP), 1 mM sodium pyruvate (Sigma-Aldrich, S8636), 1% penicillin-streptomycin (Corning, 30-002-CI), 50 μg/ml gentamicin (Corning, 30-005- CR) and 2% MEM amino acids (Sigma-Aldrich, M5550) at 37°C and 10% CO_2_. The culture medium was replaced every 3 days. To synchronize VSMCs to the G0 phase of the cell cycle, VSMCs at 70-80% confluency were incubated in serum-free DMEM containing 1 mg/ml bovine serum albumin (BSA; Sigma-Aldrich, A1933) for 48 hours. Cells were detached with 0.05% trypsin-EDTA (GIBCO, 25200056), centrifuged, resuspended, and subsequently seeded onto sterilized scaffolds in DMEM containing 10% FBS for 24 hours before downstream assays.

### 4.7. Immunofluorescence

Scaffolds were first hydrated in PBS and sterilized under UV light for 30 min prior to cell seeding. Serum-starved human VSMCs were seeded onto sterilized scaffolds and cultured in DMEM containing 10% FBS for 24 hours. Cells were then fixed with 4% paraformaldehyde (Thermo Fisher Scientific, J19943.K2), permeabilized with 0.1% Triton X-100 (Sigma-Aldrich, X100), and blocked with 3% BSA. Samples were incubated with primary antibodies against survivin (Cell Signaling, 2808) and YAP (Thermo Fisher Scientific, PA146189). F-actin was stained using phalloidin (Thermo Fisher Scientific, A12379), and nuclei were counterstained with DAPI (Thermo Fisher Scientific, P36935). Fluorescence images were acquired using a Leica DM6000 widefield microscope. Cell morphology parameters and nuclear-to-cytoplasmic intensity ratios of YAP and survivin were quantified using ImageJ (NIH).

### 4.8. RNA-sequencing

RNA-sequencing was performed to evaluate global transcriptional responses of VSMCs cultured on soft or stiff scaffolds for 24 hours. Passage 2-3 human VSMCs were seeded onto soft or stiff scaffolds at a density of 2 × 10^5^ per scaffold (24 mm × 55 mm) and cultured for 24 hours. Following culture, the cells were lysed in TRIzol (Thermo Fisher Scientific, 15596026), and total RNA was purified using the Direct-zol RNA MicroPrep kit (Zymo Research, R2062) according to the manufacturer’s instructions. RNA quality was assessed using a NanoDrop Spectrophotometer (Thermo Fisher Scientific). RNA-Seq library preparation (rRNA depletion) and sequencing (150 bp paired-end) were performed at Genewiz (South Plainfield, New Jersey)^58,59^. Raw sequence reads were processed to remove adapter sequences and low-quality bases using Trimmomatic v.0.36. The reads were then mapped to the *Homo sapiens GRCh38* reference genome (ENSEMBL) using the STAR aligner v.2.5.2b. Using featureCounts from the Subread package v.1.5.2, unique gene hit counts were calculated to quantify gene expression.

Differential expression analysis was performed using DESeq2, and data visualization was conducted using the ggplot2 package in R^60^. Differentially expressed genes (DEGs) in stiff versus soft conditions were defined using fold change > 2.0 and p-value < 0.5 thresholds. For network and clustering analyses, significantly upregulated DEGs were analyzed using STRING and Cytoscape to identify protein-protein interaction networks. Genes were grouped into four clusters using K-means clustering, and functional enrichment analysis was performed for each cluster to identify the top biological processes.

### 4.9. Statistical Analysis

All quantitative data are presented as mean ± standard deviation (SD). Statistical analyses were performed using GraphPad Prism 10. Comparisons between two groups were conducted using an unpaired two-tailed Student’s t-test. For experiments with more than two groups, one-way analysis of variance (ANOVA) was performed, followed by appropriate post hoc multiple-comparisons testing. A p-value<0.05 was considered statistically significant.

## ACKNOWLEDGEMENTS

This research was supported by grants from the National Institutes of Health (R01 HL163168, R01 AR079224, P50 AR080581).

## SUPPLEMENTAL INFORMATION

**Supplementary Figure 1:**
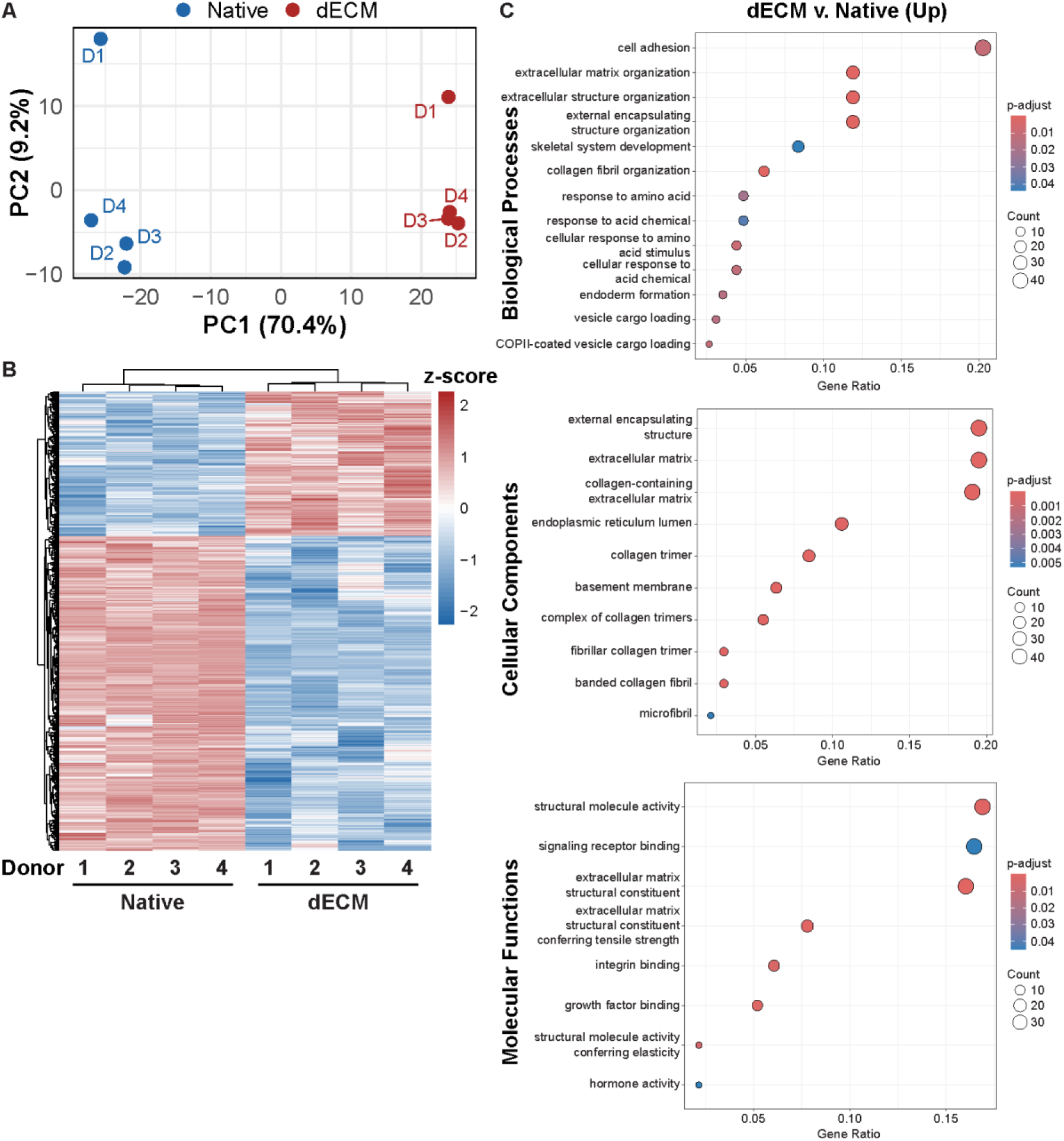
Proteomic profiling distinguishes native and dECM samples and reveals enrichment of ECM-associated functional signatures following decellularization. **(A)** Principal component analysis (PCA) of proteomic profiles from native and dECM samples. **(B)** Heatmap of selected proteins detected across native and dECM groups. **(C)** Gene ontology enrichment analysis of proteins upregulated in dECM relative to native samples, categorized by biological processes, cellular components, and molecular functions. Differential abundance analysis was performed using limma, with significance defined as p < 0.05. (n = 4 biological replicates per group).

**Supplementary Figure 2:**
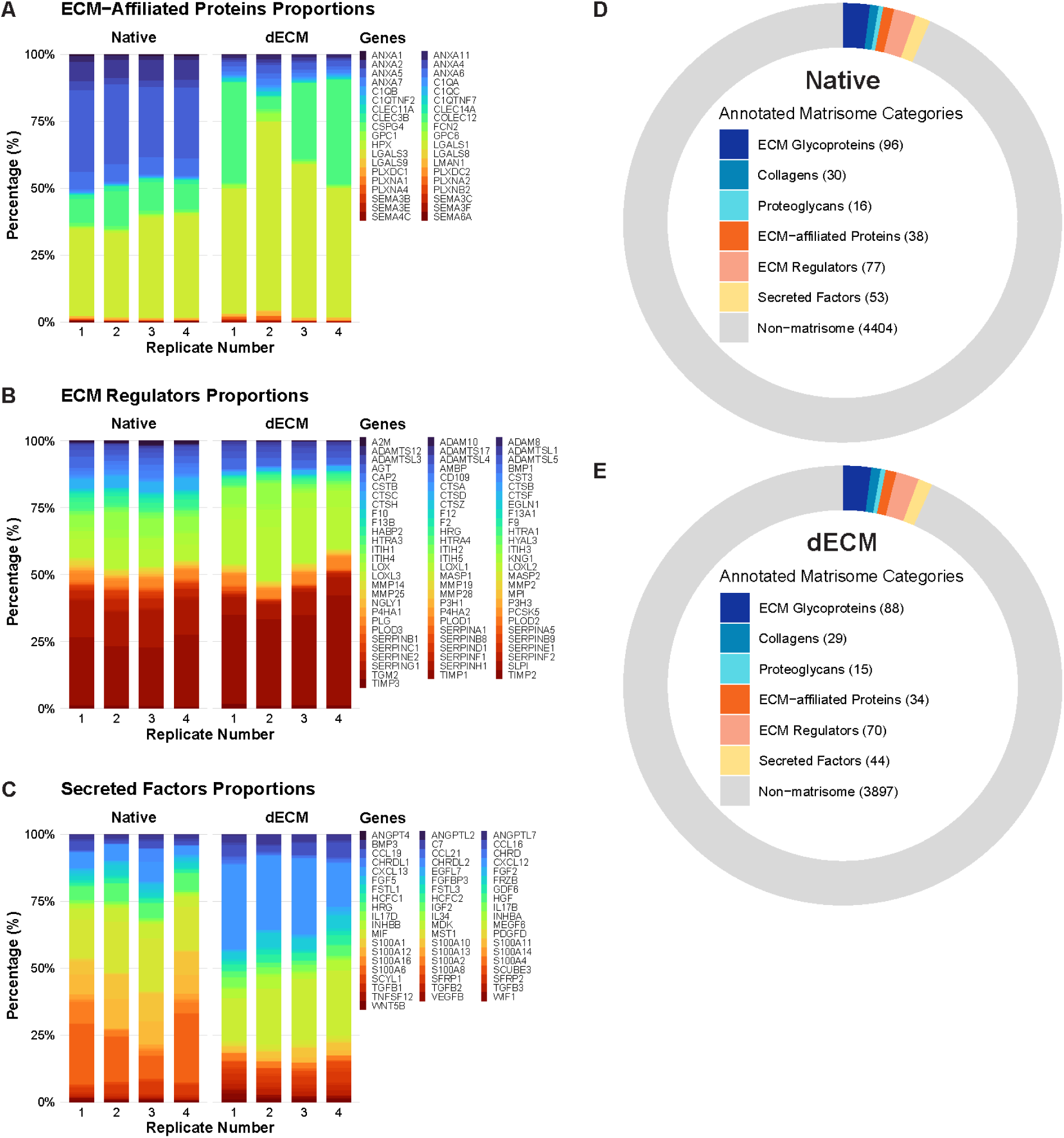
Matrisome-associated protein composition is broadly retained following decellularization. Relative protein proportions of matrisome-associated classes across native and dECM replicates, including **(A)** ECM-affiliated proteins, **(B)** ECM regulators, and **(C**) secreted factors. Distribution of annotated matrisome categories in **(D)** native and **(E)** dECM samples. Matrisome annotation was performed using MatrisomeAnalyzeR (n = 4 biological replicates per group).

**Supplementary Figure 3:**
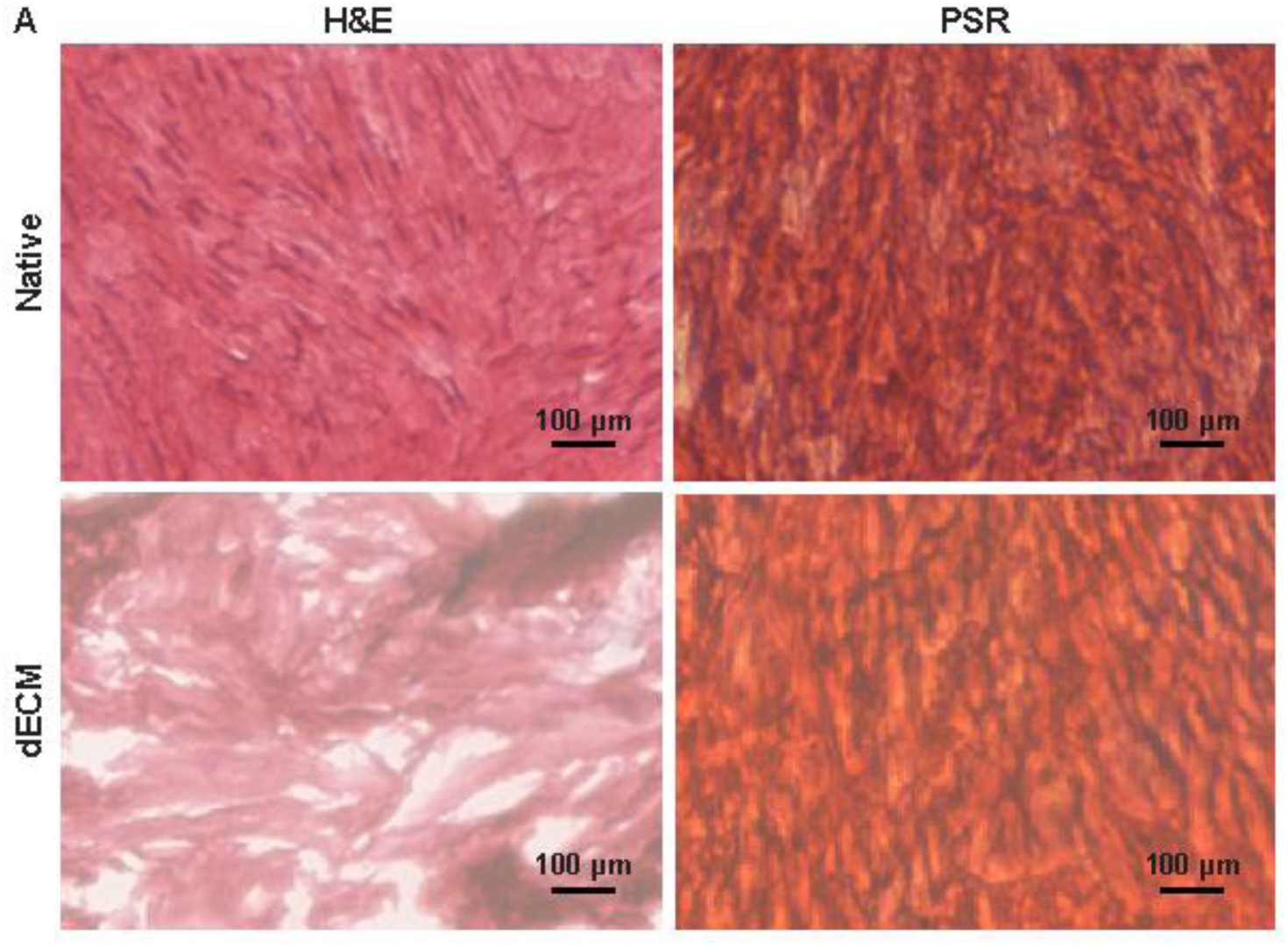
Histological staining confirms effective cellular removal and preservation of collagen-rich matrix architecture following decellularization. **(A)** Representative hematoxylin and eosin (H&E) and picrosirius red (PSR) staining of native porcine aorta and dECM samples (scale bars = 100 µm). H&E staining demonstrates marked reduction of cellular content following decellularization, while PSR staining indicates preservation of collagen-rich matrix architecture.

**Supplementary Figure 4:**
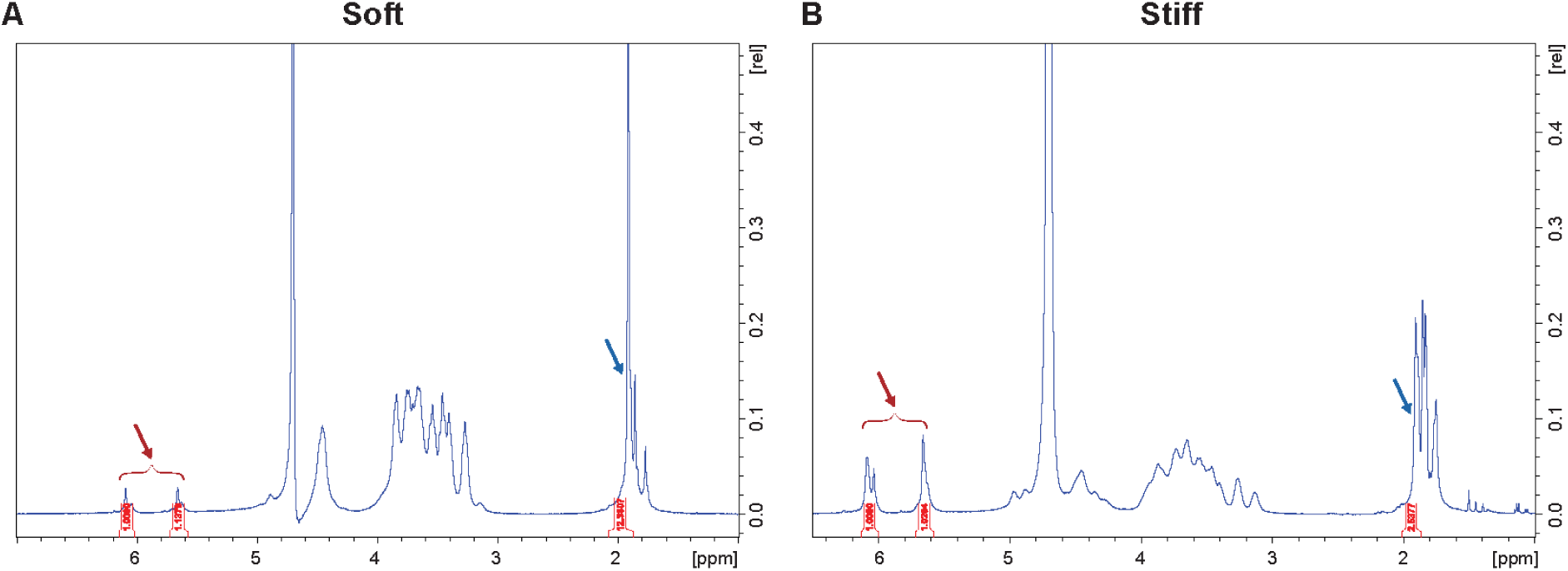
¹H NMR characterization confirms differential methacrylation of soft and stiff methacrylated hyaluronic acid (MeHA). Representative ¹H NMR spectra of methacrylated hyaluronic acid (MeHA) dissolved in D₂O for **(A)** soft MeHA and **(B)** stiff MeHA formulations. stiff formulations. The degree of methacrylation was calculated from the ratio of the integrated areas of the characteristic methacrylate proton peaks at 5.6 and 6.1 ppm (red arrows) to the N-acetyl glucosamine peak at 1.9 ppm (blue arrow). Soft MeHA exhibited a degree of methacrylation of ∼25%, whereas stiff MeHA exhibited a degree of methacrylation of ∼119%.

**Supplementary Figure 5:**
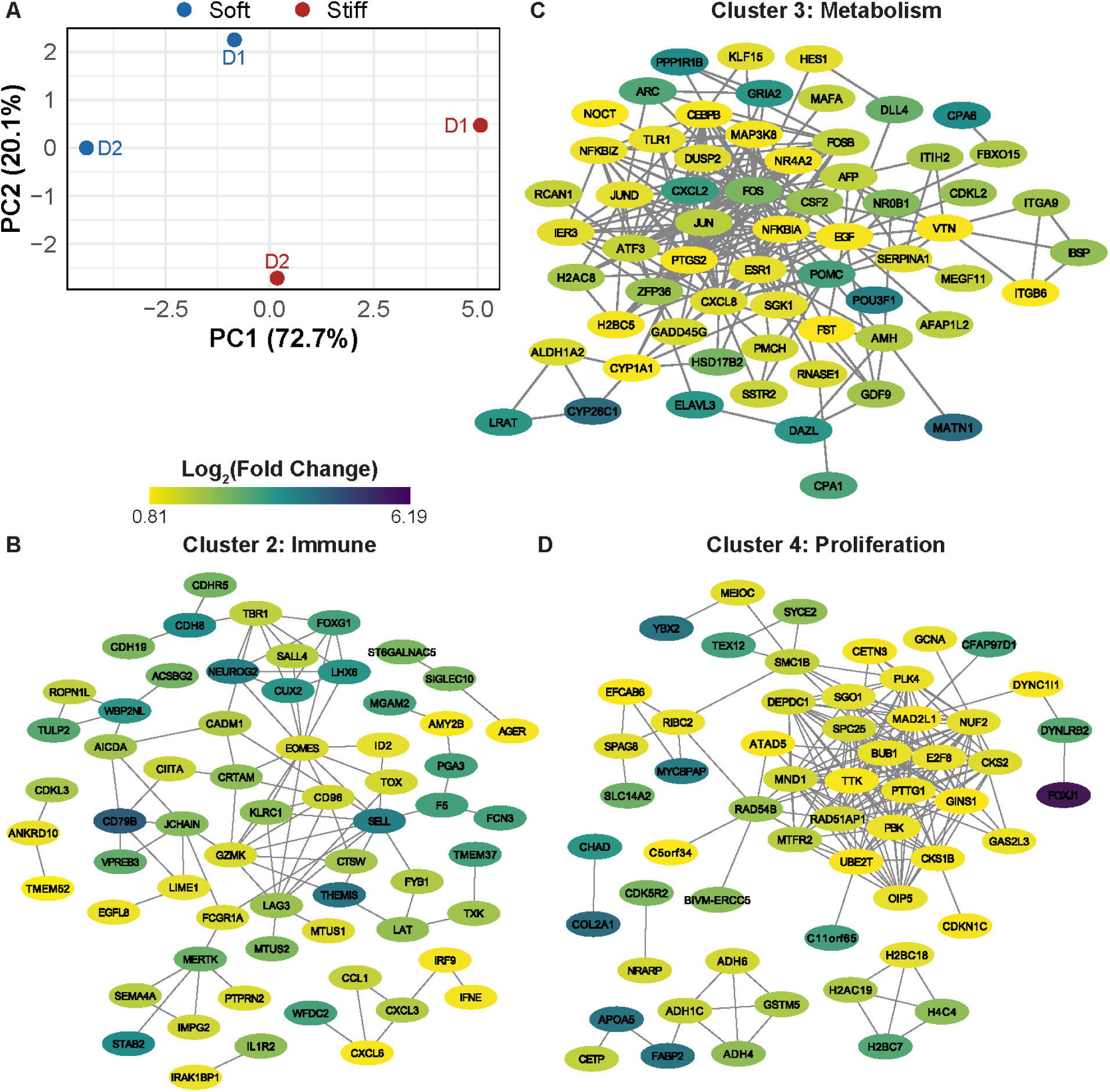
RNA-sequencing analysis identifies stiffness-responsive VSMC transcriptional programs and interaction networks. **(A)** Principal component analysis (PCA) of RNA-sequencing profiles from VSMCs cultured on soft and stiff dECM–MeHA scaffolds for 24 hours. STRING-based protein–protein interaction networks of upregulated differentially expressed genes (DEGs), grouped into clusters associated with **(B)** immune-related signaling, **(C)** metabolism/stress responses, and **(D)** proliferation. Node color indicates log2(fold change) in stiff versus soft scaffolds. RNA sequencing was performed using VSMCs from n = 2 biological donors cultured on soft and stiff scaffolds for 24 hours. Differential expression analysis was performed using DESeq2, with significance defined as p < 0.05 and fold change > 2.0. STRING clustering was performed using K-means clustering (k = 4).

**Supplementary Figure 6:**
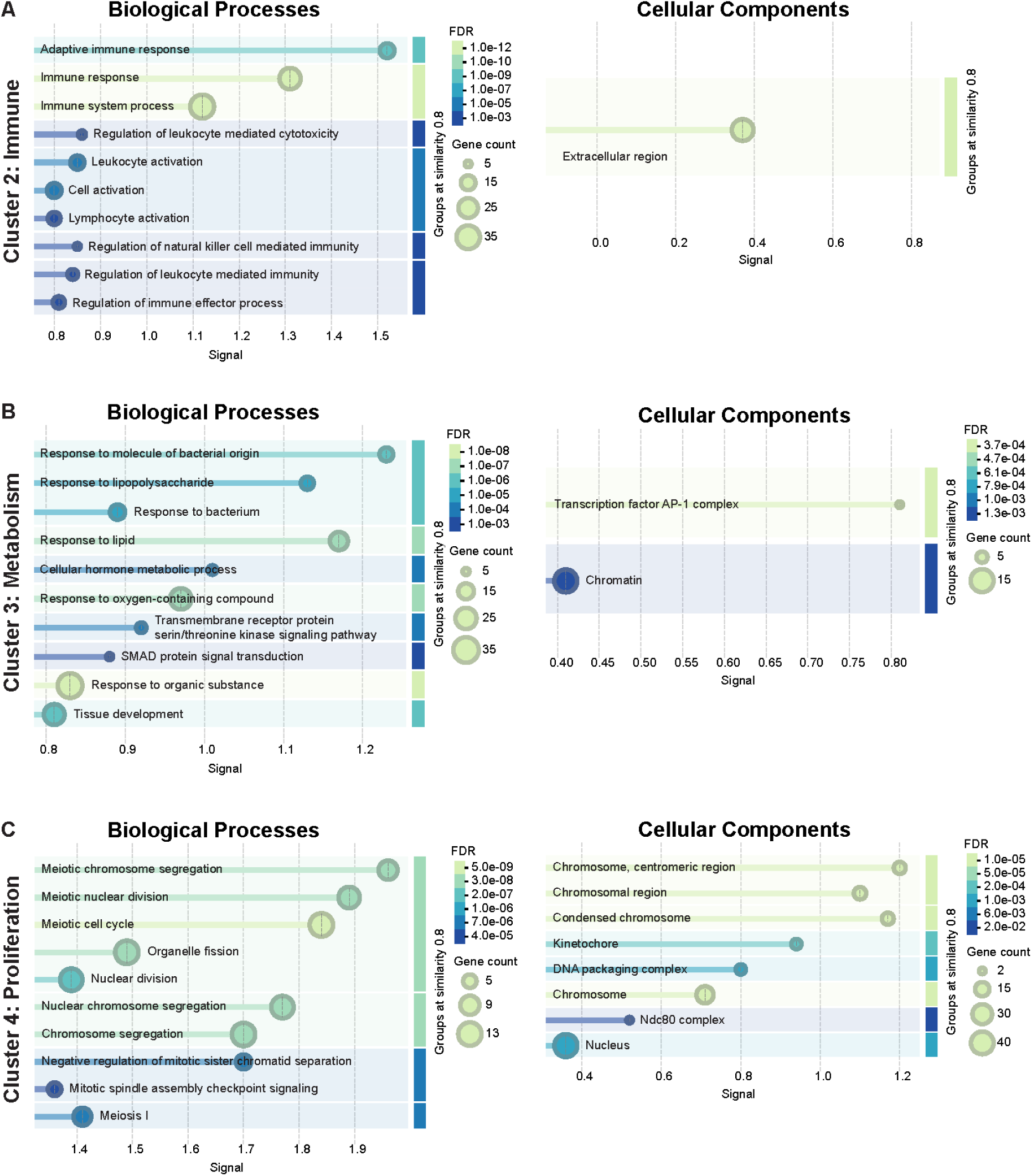
Gene ontology enrichment identifies immune-, stress-, and proliferation-associated programs within stiffness-responsive VSMC interaction clusters. Gene ontology enrichment analysis of clusters derived from upregulated differentially expressed genes (DEGs) in VSMCs cultured on stiff scaffolds. Enriched biological process and cellular component terms are shown for clusters associated with **(A)** immune-related signaling, **(B)** metabolism/stress responses, and **(C)** proliferation.

## REFERENCES

1. Diseases I of M (US) C on a NSS for C and SC. Cardiovascular Disease. In: A Nationwide Framework for Surveillance of Cardiovascular and Chronic Lung Diseases. National Academies Press (US); 2011. Accessed April 21, 2026. https://www.ncbi.nlm.nih.gov/books/NBK83160/

2. Mensah GA, Roth GA, Fuster V. The Global Burden of Cardiovascular Diseases and Risk Factors: 2020 and Beyond. J Am Coll Cardiol. 2019;74(20):2529–2532. doi:10.1016/j.jacc.2019.10.009

3. 2026 Heart Disease and Stroke Statistics: A Report of US and Global Data From the American Heart Association. doi:10.1161/CIR.0000000000001412

4. Lacolley P, Regnault V, Segers P, Laurent S. Vascular Smooth Muscle Cells and Arterial Stiffening: Relevance in Development, Aging, and Disease. Physiol Rev. 2017;97(4):1555–1617. doi:10.1152/physrev.00003.2017

5. Sehgel NL, Vatner SF, Meininger GA. “Smooth Muscle Cell Stiffness Syndrome”—Revisiting the Structural Basis of Arterial Stiffness. Front Physiol. 2015;6:335. doi:10.3389/fphys.2015.00335

6. Van den Bergh G, Opdebeeck B, D’Haese PC, Verhulst A. The Vicious Cycle of Arterial Stiffness and Arterial Media Calcification. Trends Mol Med. 2019;25(12):1133–1146. doi:10.1016/j.molmed.2019.08.006

7. Di Nubila A, Dilella G, Simone R, Barbieri SS. Vascular Extracellular Matrix in Atherosclerosis. Int J Mol Sci. 2024;25(22):12017. doi:10.3390/ijms252212017

8. Bennett MR, Sinha S, Owens GK. Vascular smooth muscle cells in atherosclerosis. Circ Res. 2016;118(4):692–702. doi:10.1161/CIRCRESAHA.115.306361

9. Zhang F, Guo X, Xia Y, Mao L. An update on the phenotypic switching of vascular smooth muscle cells in the pathogenesis of atherosclerosis. Cell Mol Life Sci CMLS. 2021;79(1):6. doi:10.1007/s00018-021-04079-z

10. Sazonova OV, Isenberg BC, Herrmann J, et al. Extracellular matrix presentation modulates vascular smooth muscle cell mechanotransduction. Matrix Biol. 2015;41:36–43. doi:10.1016/j.matbio.2014.11.001

11. Talwar S, Kant A, Xu T, Shenoy VB, Assoian RK. Mechanosensitive smooth muscle cell phenotypic plasticity emerging from a null state and the balance between Rac and Rho. Cell Rep. 2021;35(3). doi:10.1016/j.celrep.2021.109019

12. Dupont S, Morsut L, Aragona M, et al. Role of YAP/TAZ in mechanotransduction. Nature. 2011;474(7350):179–183. doi:10.1038/nature10137

13. Wheatley SP, Altieri DC. Survivin at a glance. J Cell Sci. 2019;132(7):jcs223826. doi:10.1242/jcs.223826

14. Biber JC, Sullivan A, Brazzo JA III, et al. Survivin as a mediator of stiffness-induced cell cycle progression and proliferation of vascular smooth muscle cells. APL Bioeng. 2023;7(4):046108. doi:10.1063/5.0150532

15. Krajnik A, Nimmer E, Brazzo JA, et al. Survivin regulates intracellular stiffness and extracellular matrix production in vascular smooth muscle cells. APL Bioeng. 2023;7(4):046104. doi:10.1063/5.0157549

16. Mousso T, Rice K, Tumenbayar BI, et al. Survivin modulates stiffness-induced vascular smooth muscle cell motility. APL Bioeng. 2025;9(2):026120. doi:10.1063/5.0252766

17. Blanc-Brude OP, Yu J, Simosa H, Conte MS, Sessa WC, Altieri DC. Inhibitor of apoptosis protein survivin regulates vascular injury. Nat Med. 2002;8(9):987–994. doi:10.1038/nm750

18. Simosa HF, Wang G, Sui X, et al. Survivin expression is up-regulated in vascular injury and identifies a distinct cellular phenotype. J Vasc Surg. 2005;41(4):682–690. doi:10.1016/j.jvs.2005.01.006

19. Blache U, Ford EM, Ha B, et al. Engineered hydrogels for mechanobiology. Nat Rev Methods Primer. 2022;2:98. doi:10.1038/s43586-022-00179-7

20. Geckil H, Xu F, Zhang X, Moon S, Demirci U. Engineering hydrogels as extracellular matrix mimics. Nanomed. 2010;5(3):469–484. doi:10.2217/nnm.10.12

21. Moroni F, Mirabella T. Decellularized matrices for cardiovascular tissue engineering. Am J Stem Cells. 2014;3(1):1–20.

22. Pierre V, Wu DH, Liu C, Ertugral E, Kothapalli C, Senyo SE. Tunable methacrylated decellularized heart matrix: a versatile scaffold for cardiac tissue engineering. Front Bioeng Biotechnol. 2025;13. doi:10.3389/fbioe.2025.1579246

23. Klein EA, Yin L, Kothapalli D, et al. Cell-Cycle Control by Physiological Matrix Elasticity and In Vivo Tissue Stiffening. Curr Biol. 2009;19(18):1511–1518. doi:10.1016/j.cub.2009.07.069

24. Nardone G, Oliver-De La Cruz J, Vrbsky J, et al. YAP regulates cell mechanics by controlling focal adhesion assembly. Nat Commun. 2017;8:15321. doi:10.1038/ncomms15321

25. Rickel AP, Sanyour HJ, Leyda NA, Hong Z. Extracellular Matrix Proteins and Substrate Stiffness Synergistically Regulate Vascular Smooth Muscle Cell Migration and Cortical Cytoskeleton Organization. ACS Appl Bio Mater. 2020;3(4):2360–2369. doi:10.1021/acsabm.0c00100

26. Halder G, Dupont S, Piccolo S. Transduction of mechanical and cytoskeletal cues by YAP and TAZ. Nat Rev Mol Cell Biol. 2012;13(9):591–600. doi:10.1038/nrm3416

27. Helin K. Regulation of cell proliferation by the E2F transcription factors. Curr Opin Genet Dev. 1998;8(1):28–35. doi:10.1016/s0959-437x(98)80058-0

28. Hill MA, Meininger GA. Arteriolar Vascular Smooth Muscle Cells: Mechanotransducers in a complex environment. Int J Biochem Cell Biol. 2012;44(9):1505–1510. doi:10.1016/j.biocel.2012.05.021

29. Brozovich FV, Nicholson CJ, Degen CV, Gao YZ, Aggarwal M, Morgan KG. Mechanisms of Vascular Smooth Muscle Contraction and the Basis for Pharmacologic Treatment of Smooth Muscle Disorders. Pharmacol Rev. 2016;68(2):476–532. doi:10.1124/pr.115.010652

30. Lim S, Park S. Role of vascular smooth muscle cell in the inflammation of atherosclerosis. BMB Rep. 2014;47(1):1–7. doi:10.5483/BMBRep.2014.47.1.285

31. Turner CT. Pro-inflammatory granzyme K contributes extracellularly to disease. Front Immunol. 2025;16:1620670. doi:10.3389/fimmu.2025.1620670

32. Lan F, Li J, Miao W, et al. GZMK-expressing CD8+ T cells promote recurrent airway inflammatory diseases. Nature. 2025;638(8050):490–498. doi:10.1038/s41586-024-08395-9

33. Zheng S, Yang B, Li L, et al. CRTAM promotes antitumor immune response in triple negative breast cancer by enhancing CD8+ T cell infiltration. Int Immunopharmacol. 2024;129:111625. doi:10.1016/j.intimp.2024.111625

34. Takeuchi A, Badr MESG, Miyauchi K, et al. CRTAM determines the CD4+ cytotoxic T lymphocyte lineage. J Exp Med. 2015;213(1):123–138. doi:10.1084/jem.20150519

35. Hu C, Huang W, Xiong N, Liu X. SP1-mediated transcriptional activation of PTTG1 regulates the migration and phenotypic switching of aortic vascular smooth muscle cells in aortic dissection through MAPK signaling. Arch Biochem Biophys. 2021;711:109007. doi:10.1016/j.abb.2021.109007

36. McCleland ML, Kallio MJ, Barrett-Wilt GA, et al. The Vertebrate Ndc80 Complex Contains Spc24 and Spc25 Homologs, which Are Required to Establish and Maintain Kinetochore-Microtubule Attachment. Curr Biol. 2004;14(2):131–137. doi:10.1016/j.cub.2003.12.058

37. Stratford JK, Yan F, Hill RA, et al. Genetic and pharmacological inhibition of TTK impairs pancreatic cancer cell line growth by inducing lethal chromosomal instability. PLOS ONE. 2017;12(4):e0174863. doi:10.1371/journal.pone.0174863

38. Ciccone G, Salmeron-Sanchez M. Tuning the matrix: Recent advances in mechanobiology unveiled through polyacrylamide hydrogels. Curr Opin Biomed Eng. 2025;35:100604. doi:10.1016/j.cobme.2025.100604

39. Scott KE, Fraley SI, Rangamani P. A spatial model of YAP/TAZ signaling reveals how stiffness, dimensionality, and shape contribute to emergent outcomes. Proc Natl Acad Sci. 2021;118(20):e2021571118. doi:10.1073/pnas.2021571118

40. Dong DL, Jin GZ. YAP and ECM Stiffness: Key Drivers of Adipocyte Differentiation and Lipid Accumulation. Cells. 2024;13(22):1905. doi:10.3390/cells13221905

41. Cao X, Pfaff SL, Gage FH. YAP regulates neural progenitor cell number via the TEA domain transcription factor. Genes Dev. 2008;22(23):3320–3334. doi:10.1101/gad.1726608

42. Shen Z, Stanger BZ. YAP Regulates S-Phase Entry in Endothelial Cells. PLoS ONE. 2015;10(1):e0117522. doi:10.1371/journal.pone.0117522

43. Ehmer U, Sage J. Control of proliferation and cancer growth by the Hippo signaling pathway. Mol Cancer Res MCR. 2016;14(2):127–140. doi:10.1158/1541-7786.MCR-15-0305

44. McMurtry MS, Archer SL, Altieri DC, et al. Gene therapy targeting survivin selectively induces pulmonary vascular apoptosis and reverses pulmonary arterial hypertension. J Clin Invest. 2005;115(6):1479–1491. doi:10.1172/JCI23203

45. Fukuda S, Pelus LM. Survivin, a cancer target with an emerging role in normal adult tissues. Mol Cancer Ther. 2006;5(5):1087–1098. doi:10.1158/1535-7163.MCT-05-0375

46. Grootaert MOJ, Bennett MR. Vascular smooth muscle cells in atherosclerosis: time for a re-assessment. Cardiovasc Res. 2021;117(11):2326–2339. doi:10.1093/cvr/cvab046

47. Wang M, Monticone RE, McGraw KR. Proinflammation, profibrosis, and arterial aging. Aging Med. 2020;3(3):159-168. doi:10.1002/agm2.12099

48. Fu Z, Yang S, Chang X, Liu P, Wang Y. Vascular smooth muscle cell metabolic reprogramming and phenotypic remodeling in atherosclerosis. Cell Death Discov. 2025;12(1):64. doi:10.1038/s41420-025-02932-9

49. Hao N, Yong H, Zhang F, et al. Aortic calcification accelerates cardiac dysfunction via inducing apoptosis of cardiomyocytes. Int J Med Sci. 2024;21(2):306–318. doi:10.7150/ijms.90324

50. Lee SH, Li Z, Zhang EY, et al. Precision repair of zone-specific meniscal injuries using a tunable extracellular matrix-based hydrogel system. Bioact Mater. 2025;48:400–413. doi:10.1016/j.bioactmat.2025.02.013

51. Ritchie ME, Phipson B, Wu D, et al. limma powers differential expression analyses for RNA-sequencing and microarray studies. Nucleic Acids Res. 2015;43(7):e47. doi:10.1093/nar/gkv007

52. Petrov PB, Considine JM, Izzi V, Naba A. Matrisome AnalyzeR – a suite of tools to annotate and quantify ECM molecules in big datasets across organisms. J Cell Sci. 2023;136(17):jcs261255. doi:10.1242/jcs.261255

53. Burdick JA, Chung C, Jia X, Randolph MA, Langer R. Controlled Degradation and Mechanical Behavior of Photopolymerized Hyaluronic Acid Networks. Biomacromolecules. 2005;6(1):386–391. doi:10.1021/bm049508a

54. Seidlits SK, Khaing ZZ, Petersen RR, et al. The effects of hyaluronic acid hydrogels with tunable mechanical properties on neural progenitor cell differentiation. Biomaterials. 2010;31(14):3930–3940. doi:10.1016/j.biomaterials.2010.01.125

55. Khetan S, Guvendiren M, Legant WR, Cohen DM, Chen CS, Burdick JA. Degradation-mediated cellular traction directs stem cell fate in covalently crosslinked three-dimensional hydrogels. Nat Mater. 2013;12(5):458–465. doi:10.1038/nmat3586

56. Tsanaktsidou E, Kammona O, Labude N, et al. Biomimetic Cell-Laden MeHA Hydrogels for the Regeneration of Cartilage Tissue. Polymers. 2020;12(7):1598. doi:10.3390/polym12071598

57. Hossain Rakin R, Kumar H, Rajeev A, et al. Tunable metacrylated hyaluronic acid-based hybrid bioinks for stereolithography 3D bioprinting. Biofabrication. 2021;13(4):044109. doi:10.1088/1758-5090/ac25cb

58. Zhang EY, Blanch TE, Ahmed SB, Jiang X, Dyment NA, Heo SC. Epigenetic regulation and mechanobiological adaptation in tenocytes during maturation. APL Bioeng. 2025;9(2):026127. doi:10.1063/5.0271050

59. Zhang Y, Zhang EY, Cheung C, et al. Epigenetic dynamics in meniscus cell migration and its zonal dependency in response to inflammatory conditions. APL Bioeng. 2025;9(1):016109. doi:10.1063/5.0239035

60. Love MI, Huber W, Anders S. Moderated estimation of fold change and dispersion for RNA-seq data with DESeq2. Genome Biol. 2014;15(12):550. doi:10.1186/s13059-014-0550-8

